# A Simple Deep Learning Approach for Detecting Duplications and Deletions in Next-Generation Sequencing Data

**DOI:** 10.1101/657361

**Authors:** Tom Hill, Robert L. Unckless

## Abstract

Copy number variants (CNV) are associated with phenotypic variation in several species. However, properly detecting changes in copy numbers of sequences remains a difficult problem, especially in lower quality or lower coverage next-generation sequencing data. Here, inspired by recent applications of machine learning in genomics, we describe a method to detect duplications and deletions in short-read sequencing data. In low coverage data, machine learning appears to be more powerful in the detection of CNVs than the gold-standard methods or coverage estimation alone, and of equal power in high coverage data. We also demonstrate how replicating training sets allows a more precise detection of CNVs, even identifying novel CNVs in two genomes previously surveyed thoroughly for CNVs using long read data.

Available at: https://github.com/tomh1lll/dudeml

## Introduction

Copy number variation (CNV) of DNA sequences is responsible for functional phenotypic variation in many organisms, particularly when it comes to causing or fighting diseases (Sturtevant 1937; Inoue and Lupski 2002; Rastogi and Liberles 2005; Jennifer L. Newman. 2006; Redon *et al.* 2006; Unckless *et al.* 2016). Despite its importance, properly detecting copy number variants is difficult and so the extent that CNVs contribute to phenotypic variation has yet to be fully ascertained (Redon *et al.* 2006; Chakraborty *et al.* 2017). This detection difficulty is due to challenges in aligning CNVs, with similar copies being combined in both Sanger-sequencing and with mapping short-read NGS data to a reference genome lacking the duplication (Redon *et al.* 2006; Ye *et al.* 2009). Several tools have been developed to detect these CNVs in next-generation sequencing (NGS) data, but for proper accuracy, they require high coverages of samples (for the detection of split-mapped reads, or better estimations of relative coverage), long-reads (able to bridge the CNVs) or computationally intensive methods (Redon *et al.* 2006; Ye *et al.* 2009; Chen *et al.* 2016; Chakraborty *et al.* 2017). This limits the ability to detect CNVs between samples sequenced to relatively low coverages, with short reads on lower quality genomes.

The recent development of numerous machine learning techniques in several aspects of genomics suggests a role for machine learning in the detection of copy number variants (Rosenberg *et al.* 2002; Sheehan and Song 2016; Schrider *et al.* 2017; Schrider and Kern 2018). Contemporary machine learning methods are able to classify windows across the genome with surprising accuracy, even using lower quality data (Kern and Schrider 2018). Additionally, machine learning techniques are generally less computationally intensive than other modern methods such as Approximate Bayesian computation, because the user providing a training set for the supervised detection of classes (Beaumont *et al.* 2002; Schrider and Kern 2018).

Here we introduce a novel deep-learning-based method for detecting duplications and deletions, named ‘**Du**plication and **De**letion Classifier using **M**achine **L**earning’ (dudeML). We outline our rationale for the statistics used to detect CNVs and the method employed, in which we calculate relative coverage changes across a genomic window (divided into sub windows) which allows for the classification of window coverages using different machine learning classifiers. Using both simulated and known copy number variants, we show how dudeML can correctly detect copy number variants and outperforms basic coverage estimates alone.

## Methods

### Machine learning method and optimization

Inspired by recent progress in machine learning for population genomics (Schrider and Kern 2016; Kern and Schrider 2018; Schrider and Kern 2018), we sought to develop a method to accurately and quickly classify the presence or absence of copy number variants in genomic windows using a supervised machine learning classifier. Based on previous software and methods for copy number detection (Ye *et al.* 2009; Chen *et al.* 2016), we identified a number of statistics that may help determine if a duplication or deletion is present in a particular window. We reasoned that both standardized and normalized median coverage should indicate if a window is an outlier from the coverage (Figure 1, black), and that the standard deviation increases in regions with higher coverage, decreases in regions with lower coverage but increase dramatically at CNV edges due to rapid shifts in coverage (Figure 1, grey). Another component of some CNV detection algorithms are unidirectional split mapped reads which also indicate the breakpoint of a structural variant such as a deletion or tandem duplication (expected at the red/blue borders in Figure 1) (Ye *et al.* 2009; Palmieri *et al.* 2014).

**Figure 1.**
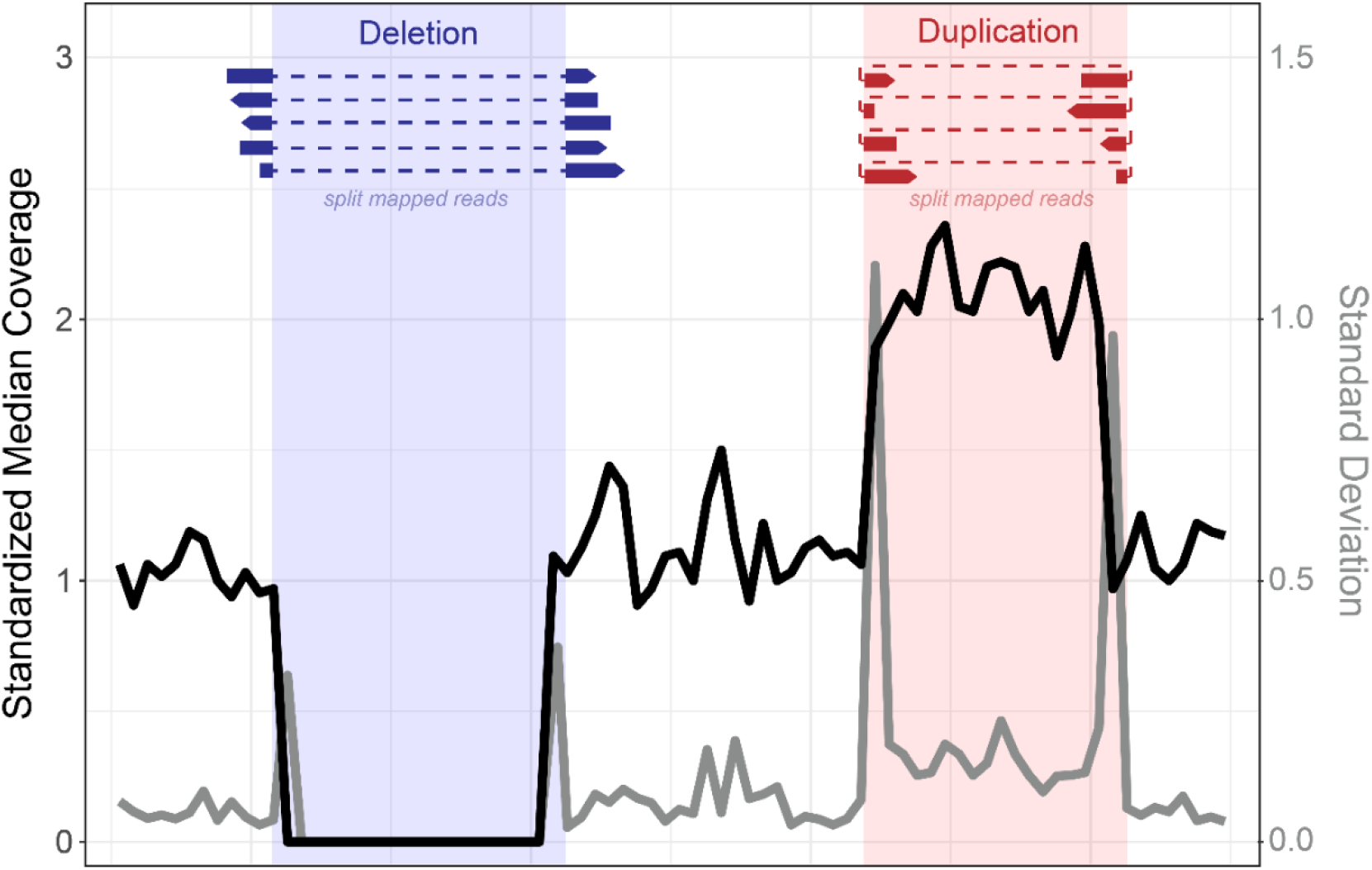
Schematic demonstrating the rationale behind each statistic used to initially determine the presence/absence of each copy number variant. We expect the Standardized median coverage (black line) to increase in duplications (red) and decrease in deletions (blue). We expect the standard deviation of the standardized coverage to greatly increase at the edges of CNVs (grey line). At the borders of CNVs we also expect an increase in split mapped reads, specifically across the edges of deletions (dark blue) or within a tandemly duplicated region (dark red).

In this classifier, we used these measures across a set of windows to define the copy number and CNV class of the focal window at the center (Figure 2A). Initially, we sought to identify which of the statistics (and in what windows) are most useful for determining the presence or absence of a copy number variant, relative to a reference genome. To do this, we simulated tandem duplications and deletions (100-5000bp) across the *Drosophila melanogaster* reference chromosome 2L. We then simulated 100bp paired-end reads for this chromosome using WGsim (Li 2012) and mapped these to the standard reference 2L using BWA and SAMtools (Li and Durbin 2009; Li *et al.* 2009), with repeats masked using RepeatMasker (Smit and Hubley 2015). We also simulated a second set of CNVs and related short read data as a test set.

**Figure 2.**
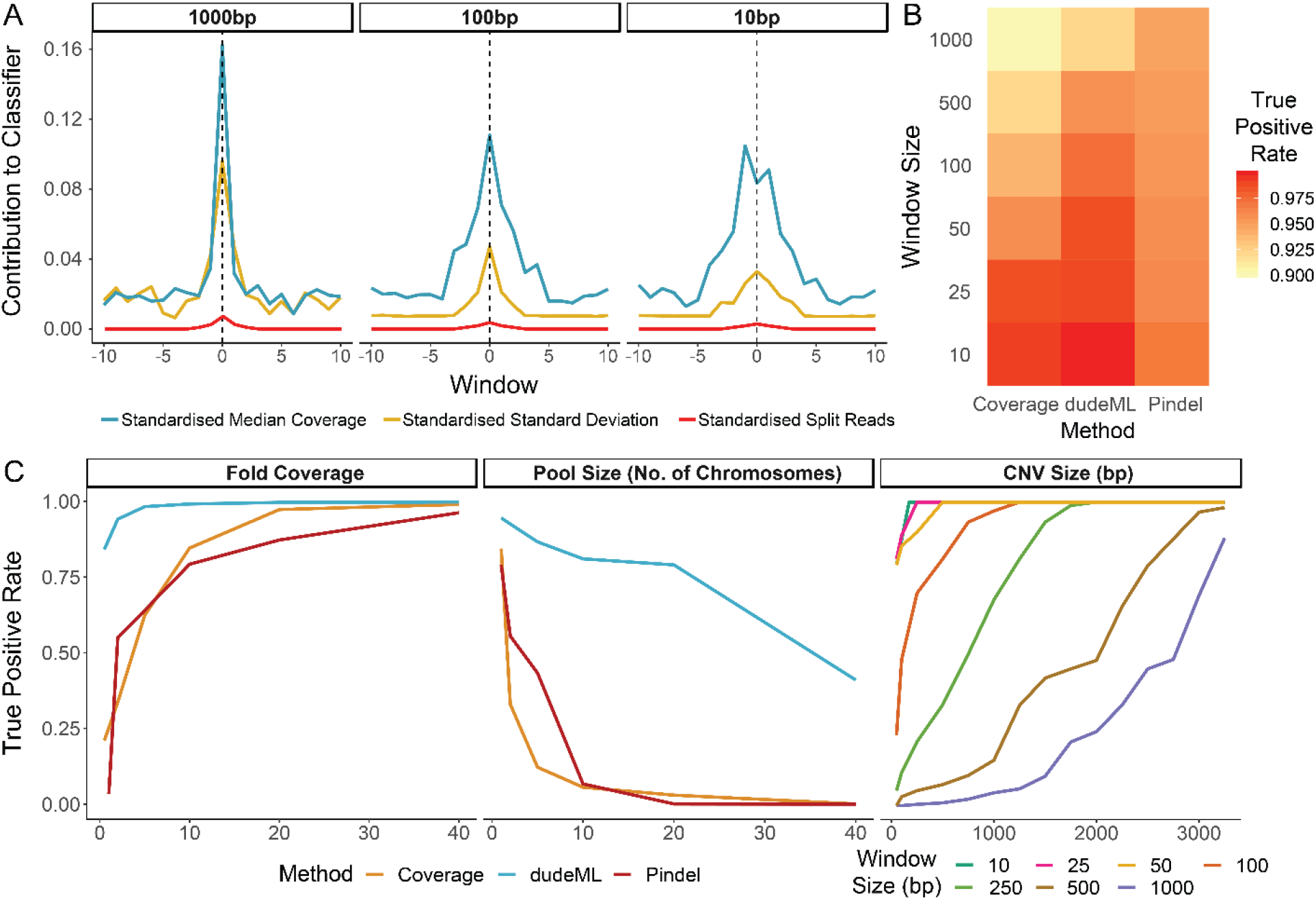
**A.** Relative contribution of each statistic to the classification of copy number variants, across windows in increasing distance from the focal window (dashed lined), separated by window size. **B.** The true positive rate of identification of simulated CNVs based on either, median coverage of the window, dudeML and Pindel. For Pindel, the overlap of called and true CNVs, rounded to the nearest window size, was used. **C.** Comparison of detection of copy number variants between Pindel, pure coverage estimations and using dudeML for varying parameters. Detection rate decreases across all methods with decreasing coverage and with increasing pool sizes. dudeML loses the ability to detect smaller CNVs with increasing window size (only shown for dudeML). Note that in **B** and **C**, for all comparisons, windows which cannot be examined in all cases (including repetitive regions) have been removed.

To identify candidate CNVs, we calculated the statistics derived above in windows between 10bp and 1000bp (sliding the same distance). We reformatted the data to vectors including the statistics for a focal sub window and 10 sub windows upstream and downstream, creating a set of statistics describing the 20 sub windows around a focal sub window, for every window set on the chromosome. We then assigned each window a class, based on the known copy number and known class (deletion, duplication or normal) for the focal sub window. We trained a random forest classifier with 100 estimators (Pedregosa *et al.* 2011) to extract what features are necessary to classify the central sub window as containing a CNV or not. We examined the contribution of statistics to classifying focal sub-windows and qualitatively removed those unimportant to the classifier e.g. statistics which appeared to not contribute to classification in any degree in any sub windows were removed upon visual inspection. This scripts and tutorial for this process are available at https://github.com/tomh1lll/dudeml, including the tool for detecting CNVs.

To further hone the method we determined how window size (10 - 1000bp), number of windows (1 - 41), coverage of data (0.2 - 40) the frequency of CNV in a pool (0.05 - 1), and how the machine learning model affects the ability to correctly classify a CNV in simulated data (Random Forest 100 estimators and 500 estimators, Extra Trees 100 and 500 estimators, Decision Tree, and Convolutional Neural Network classifiers) (Pedregosa *et al.* 2011). In each case we changed only one variable, otherwise coverage was set at 20-fold, window-size was set at 50bp, the number of sub windows each side was set to 5 and the model was set as Random Forest (100 estimators). For all comparisons (coverages, window sizes, number of windows or model comparisons) we counted the number of True and False positive CNVs and estimated a receiver operating characteristic curve (Brown and Davis 2006).

We used bedtools (Quinlan and Hall 2010) and RepeatMasker (Smit and Hubley 2015) to identify regions on chromosome 2L without high levels of repetitive content. Following this, we simulated 2000 duplications and 2000 deletions across these regions, varying in size between 100bp and 5000bp. To assess a machine learning classifiers ability to detect CNVs across pooled data, for three replicates, we created a further subset of CNVs present at different frequencies in pools of chromosomes, for pools of 2 (the equivalent of sequencing an outbred diploid individual), 5, 10 and 20 chromosomes, allowing the CNV to vary in frequency between 5% and 100% across samples, based on the number of chromosomes simulated (e.g. a 50% minimum in a pool of 2 chromosomes, equivalent to a heterozygous CNV, and a 5% minimum in a pool of 20, equivalent to a singleton CNV in a pool of 10 diploid individuals). This process was repeated twice to create independent test and training sets, both with known CNVs.

We generated chromosomes containing simulated CNVs and simulated reads for these chromosomes using WGsim (Li 2012). We simulated reads to multiple median depths of coverage per base, between 0.2 to 20. We then combined all reads for each pool set and mapped these reads to the *D. melanogaster* iso-1 reference 2L using BWA and SAMtools (Li and Durbin 2009; Li *et al.* 2009).

For each data set, of varying window sizes, coverages and pool sizes, we then reformatted each window as described above to give the statistics for the focal window and 5 windows up and downstream, unless otherwise stated. For each training set, we defined each vector by their presence in a duplication, deletion or neither. For each window we also assigned the number of copies found of that window per chromosome, e.g. 0 for a fixed deletion, 0.5 for a deletion found in 50% of chromosomes, 1.75 for a duplication found in 75% chromosomes etc. We then used SKlearn to train a classifier based on the vectors assigned to each class (Pedregosa *et al.* 2011). The classifiers were then used to assign classes to windows in the test sets, which were then compared to their known designations to identify the true positive detection rate of each set.

### Testing the classifier on real data with known CNVs

To test the classifier in known copy number variants, we downloaded the *D. melanogaster* iso-1 and A4 reference genomes (Dos Santos *et al.* 2015; Chakraborty *et al.* 2017). Then, based on (Chakraborty *et al.* 2017), we extracted windows with known duplications and deletions relative to each other, for example a tandem duplication present in one genome but not the other would appear as a deletion. We downloaded short reads for each *D. melanogaster* genome (iso-1: SRA ERR701706-11, A4: http://wfitch.bio.uci.edu/~dspr/Data/index.html) and mapped them to both genomes separately using BWA and SAMtools (Li and Durbin 2009; Li *et al.* 2009). Using the previously described methods, we calculated the coverage statistics for each window of each genome using bedtools and custom python scripts. Using the training set described previously, we then classified each window of the iso-1 and A4 strains mapped to both their own genome and the alternative reference and compared to the previously detected CNVs, this allowed us to find potential false-positives that may be due to reference genome issues.

For each dataset, we also simulated 100 independent training sets, which we used to test the effectiveness of bootstrapping the random forest classifier. Each window was reclassified for each bootstrap training set, which are then used to calculate the consensus state for each window and the proportion of boostrap replicates supporting that states.

Finally, to validate any apparent ‘False-Positive’ CNVs identified with our machine learning classifier, we downloaded Pacific Bioscience long read data for both Iso-1 and A4 (A4 PacBio SRA: SRR7874295 - SRR7874304, Iso-1 PacBio SRA: SRR1204085 - SRR1204696), and mapped this data to the opposite reference genome. For each high confidence (greater than 95% of bootstraps) ‘False-Positive’ CNV, we manually visualized the PacBio data in the integrative genomics viewer (Robinson *et al.* 2011), looking for changes in coverage and split-mapped reads. For a randomly chosen group of these CNVs, we designed primers and confirmed CNVs using PCR (Supplementary Data 1 & 2). We designed primer pairs around each CNV to assess product size differences between strains, as well as inside the CNV for strain specific amplification for deletions or laddering in the case of duplications. PCR products from primer sets in both Iso-1 and A4 were then run on a 2% gel using gel electrophoresis (Supplementary Figure 5).

## Results and Discussion

### A machine learning classifier can detect CNVs with high accuracy

We sought to develop a quick, simple and accurate classifier of copy number variants in next generation sequencing data (Pedregosa *et al.* 2011; Schrider and Kern 2016; Schrider *et al.* 2017). First, we assessed how useful multiple statistics are in the detection of non-reference duplications and deletions in short-read next-generation sequencing data (Figure 1). We simulated short read data for a chromosome containing multiple insertions and deletions relative to a reference genome and mapped these reads to the original reference chromosome. For windows across the chromosome we then calculated several statistics thought to be helpful for detecting copy number variants (CNVs) including standardized and normalized median coverage, the standard deviation of the standardized or normalized coverage within each window, and the number of split mapped reads across the window. We reasoned that each of these statistics can signal the increase or decrease of copy number of a sequence relative to a reference genome (Figure 1, see Materials and Methods). For each focal window we also included these statistics for neighboring windows. These vectors of statistics for windows with known CNVs are then fed into a machine learning classifier, which identifies the values most important to the correct classification of copy number. For simplicity we will refer to this classifier as the **Du**plication and **De**letion Classifier using **M**achine **L**earning (dudeML) moving forward. The tool developed as a wrapper for the pipeline, instructions for installation, specifics of the pipeline for detecting copy number variants, and the location of test data used in this manuscript are available at https://github.com/tomh1lll/dudeml.

Using dudeML on high coverage (>20-fold), simulated copy number variants, we find that both standardized and normalized median coverage and standard deviation are important for classifying a window. However, because normalized coverage relies on knowing the coverage distribution of a sample, we chose to remove this statistic from further analysis. Surprisingly, the number of split reads (reads where two ends map to different regions of the genome) is relatively unimportant for finding CNVs (Figure 2A). Though the breadth of a distribution will vary depending on the window-size and mean size of the CNV, the most important windows for classifying a CNV appear to be the focal window and up to 5 windows up and downstream of the focal window (Figure 2B). On a related note, increasing the number of windows surrounding the focal window decreases the true-positive rate due to a repeat content interfering with the classifier (Supplementary Figures 1 & 2, true-positive rate ~ window number, GLM t-value = −12.056, p-value = 2.478e-33). We also find different statistics have different contributions across different window sizes, for example, larger windows are more likely to include the edges of the CNV so standard deviation is more important for CNV classification in larger windows (Figure 2A). However, larger windows appear to have lower true-positive rates, again due to the increased chance of overlapping with repeat content (Supplementary Figures 1 & 2, true-positive rate ~ window size: GLM t-value = - 2.968, p-value = 0.00303).

We also compared different supervisor machine learning classifiers and found little qualitative difference between them, though the most successful classifier on simulated data was a Random Forest Classifier (Supplementary Figure 1 & 2, true-positive rate ~ classifier GLM t-value = 5.758, p-value = 8.65e-09), with no significant difference between 100 and 500 estimators (GLM t-value = −0.133, p-value = 0.246) (Pedregosa *et al.* 2011).

For this high coverage simulated data (20-fold coverage), containing known CNVs, we compared dudeML to the prediction of a CNV based on copy number alone (rounding the coverage to the nearest whole value), or Pindel (Ye *et al.* 2009), a frequently used method for deletion and duplication prediction. dudeML has a higher rate of success predicting the presence of a CNV and the windows in which the CNV starts/ends (Figure 2B, including false-positives and negatives in all cases). However, the success of a window-based approach decreases as windows increase in size, for both the machine learning classifier and coverage alone, with Pindel having a higher success rate for CNVs compared to dudeML using sub-windows greater than ~250bp (Figure 2B). As dudeML is not optimized to function in regions with repetitive content, it also lacks the ability to detect CNVs in repetitive regions, unlike Pindel (Ye *et al.* 2009). Overall, dudeML has higher success at fine window sizes or in lower coverage data (Figure 2) while for very high coverage data for large CNVs, Pindel appears to be superior (Figure 2).

### CNV machine learning classifiers are relatively agnostic to coverage and can detect CNVs in pooled data with relatively high accuracy

We next tested the extent that changing different parameters affected dudeML’s ability to correctly detect CNVs, compared to pure copy number estimates (rounding the coverage to the nearest whole value), or Pindel (Ye *et al.* 2009). We examined the effects of decreasing coverage, increasing window size and increasing the number of sub windows on correctly classifying CNVs with dudeML, in comparison to Pindel and coverage estimates for decreasing coverage. As expected, all three methods (dudeML using eleven 50bp windows, Pindel and pure coverage) have a decreasing true-positive rate with decreasing mapped coverage (Supplementary Figures 1 & 2, true-positive rate ~ coverage GLM t-value = 209.4 p-value < 2e-16). However, the correct detection of variants and their copy number is above 95% for euchromatic regions with dudeML until coverage is below 2-fold (Figure 2C, 99.8% above 10-fold, 48% at 0.5-fold). This can also be seen in the ROC curves for duplications and deletions at different sample coverages (Supplementary Figure 1) and in the proportion of true-positives found (Supplementary Figure 2). Note that the ROC curves include all windows across the genome (including windows with no CNVs), potentially inflating the true-positive rate (Supplementary Figure 1), while the second instance, CNVs in regions of the genome not analyzed are also included, inflating the false-negative rate (Supplementary Figure 2).

Compared to dudeML, Pindel and pure coverage estimation decreases in effectiveness faster than linearly (Figure 2C, >77% above 10-fold coverage, <3.5% at 0.5-fold coverage). As Pindel relies on split-mapped reads of certain mapping orientations to detect copy number variants, low coverage data likely lacks an abundance of these reads for the correct detection of CNVs (Ye *et al.* 2009). Similarly, the spurious nature of data at low coverages prevents pure relative coverage comparisons from being useful. With machine learning however, the classifier relies on thousands of similar examples in each state to more reliably predict the presence or absence of a CNV, if the training data is similar to the sampled data. In fact, correctly predicting a CNV in data of decreasing coverage with a poorly optimized training set has a similar success rate as pure-coverage alone (Supplementary Figure 3), highlighting the importance of a training set as like the true data as possible.

Often, populations are sequenced as pools of individuals instead of individually prepared samples, due to its reducing the cost of an experiment while still providing relatively high power for population genetic inference (SchlÖtterer *et al.* 2014). We simulated CNVs at varying frequencies throughout pools of chromosomes (poolseq) to assess dudeML’s ability to detect the correct number of copies of a gene in a population. We generated simulated pools as both test data and training sets of 1 (haploid or inbred), 2 (diploid, 50% coverage), 5, 10, 20 and 40 chromosomes (pools at 1-fold coverage for each chromosome), again, we compared this to Pindel’s ability to detect the CNV and relative coverage estimates. In all three cases, as the pool size increases, the ability to detect the correct number of copies of a window (or to detect copy number variants at all in Pindel) decreases (Figure 2). However, for copy number variants above ~20% frequency, dudeML is able to correctly predict their presence an average of 87% of the time, suggesting that for poolseq, dudeML has high confidence in calling CNVs compared to pure coverage of Pindel, but low confidence in accurate frequency prediction (< 21% success rate in both methods). This is likely as the changes in relative coverage and proportion of split reads becomes so slight that the proper detection is not feasible. For example, finding a fixed duplication in a single chromosome sample requires detecting a 2-fold change in coverage, while a duplication in one chromosome in a pool of 20 requires detecting a 1.05-fold change in coverage. With variance in coverage existing in even inbred samples, this makes proper CNV detection at high resolution in pools unfeasible. As before, a machine learning classifier has relatively higher success (Figure 2C), though still low, ranging from 47-94% proper detection. If the goal is, however, to detect changes in copy number variants between two samples (either over time or between two geographically distinct samples), dudeML should be enough to detect changes at around a ~20% resolution with relatively high confidence (Figure 2C), such that it may not be possible to get accurate frequency estimates in the pool, but should be able to infer the presence of duplications/deletions with at least 20% frequency, or distinguish between CNVs present at 20% frequency and 40% frequency.

### Resampling increases CNV machine learning classifier accuracy

To further tune the accuracy of our classifier, we tested its effectiveness on the detection of copy number variants in real data, as opposed to simulated copy number variants in simulated reads (though with a classifier still using simulated CNVs and simulated data for training). We therefore downloaded two *Drosophila melanogaster* reference genomes – both assembled with long-read data – with identified duplications and deletions relative to each other (A4 and Iso-1) (Chakraborty *et al.* 2017). When data from one reference is mapped to the other, regions with copy number variants show signatures of changes in standardized coverage and standard deviation as seen in simulated data (Figure 1, Supplementary Data 1).

As before we trained the classifier based on median coverage and standard deviation of simulated CNVs and standard regions, then predicted windows with duplications or deletions using a random forest approach (Pedregosa *et al.* 2011). Strangely, and unseen in simulated examples, the proportion of false-positives was extremely high, with over ten times the number of false-positives compared to true-positives (Table 1). We suspected that artefacts and false CNVs were caused by real structural variants that went undetected in the original training set and areas with inconsistent mapping rates, so we attempted to control for this by resampling across multiple training sets with independently generated CNVs. We generated 100 independent training sets across both the Iso-1 and A4 reference genomes to create 100 independent classifiers. Following this we performed a bootstrapping-like approach, predicting the copy number of each window based on each of the 100 classifiers and taking the consensus of these calls. As the number of replicates increased, the false-positive rate dropped dramatically with little effect on the true-positive rate (Table 1, Figure 3B). In fact, taking CNVs found in at least 98% of the bootstraps removed all but 17 false-positives. This did however remove some low confidence but real duplications, and therefore provides a conservative set of CNVs (Figure 3A) (Chakraborty *et al.* 2017). This suggests that multiple independent training sets can remove any artefacts found in a single training set which may lead to false calls (Table 1, Figure 3A).

**Figure 3:**
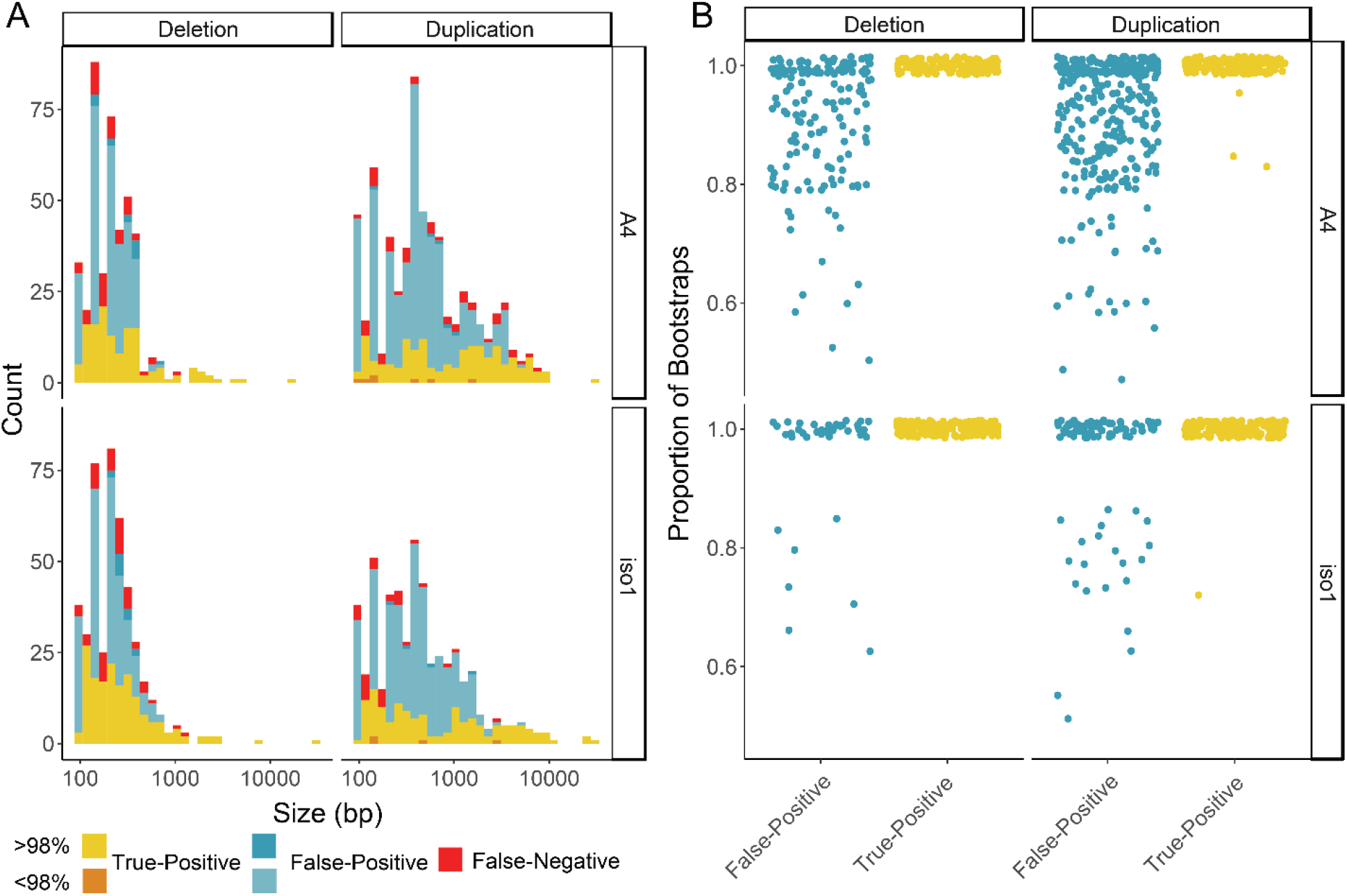
**A.** Number of CNVs detected in *Drosophila melanogaster* strains with known CNVs relative to each other after 100 bootstraps. CNVs are labelled by their previously known detection in these strains (‘True-Positive’), their lack of knowledge in these strains (‘False-Positive’) and if known CNVs were missed (‘False-Negative’). CNVs are also labelled based on the proportion of bootstraps confirming them. **B.** The proportion of bootstraps for each detected CNV in **A**, separated by if they are a false-positive, true-positive, duplication or deletion and by each strain.

**Table 1:**
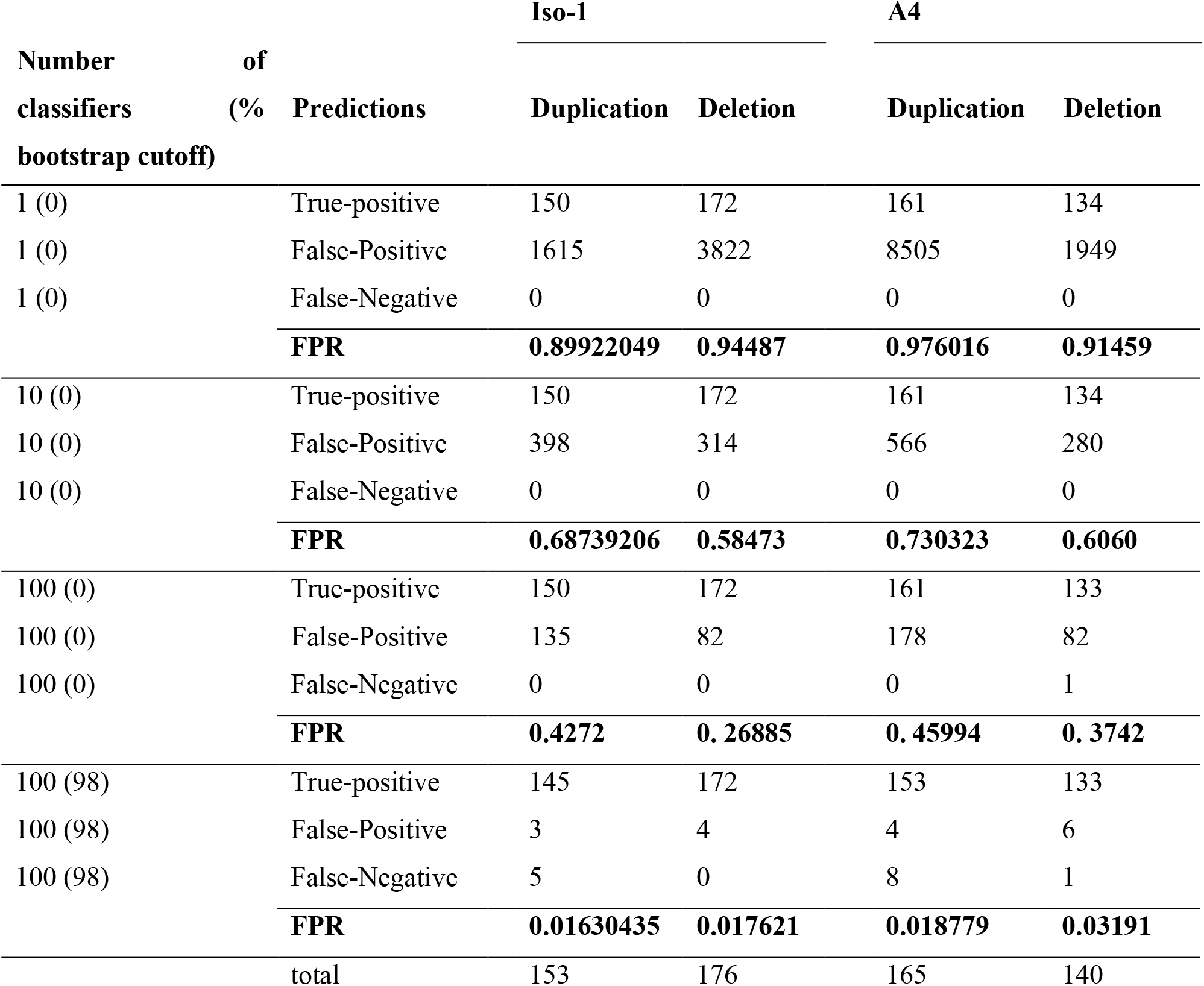
The number of predicted copy number variants in each strain (relative to the alternate strain), compared to previously identified copy-number variants (Chakraborty *et al.* 2017), across differing numbers of bootstraps and cutoffs, including the false-positive rate (FPR) for each category. Note that previously called CNVs not examined here (as they are in regions of the genome not analyzed) are included as False-Negatives in brackets for transparency.

As so many false-positives are found with high confidence across both samples, we next visually inspected the regions of the genome called as False-Positive CNVs in at least 95 of 100 bootstraps (Supplementary Figure 4, 106 duplications and 64 deletions across both strains). We extracted long reads (> 250bp) from PacBio data for both strains and mapped these to the opposite strains genome, which we then visualized in the integrative genomics viewer (Robinson *et al.* 2011). All False-positive CNVs examined show similar signatures to true-positive copy numbers (e.g. split-mapped reads across regions of 0 coverage for deletions, and supplementary alignments of reads in regions of high coverage for duplications), suggesting that they may be real CNVs and not false-positives (or at least have similar signatures to real CNVs, 18 examples given in Supplementary Data 1). We further PCR validated 12 of these CNVs, chosen at random (Supplementary Figure 5, Supplementary Data 2). While we could validate all deletions, we found no length variation in PCR product for putative duplications for primers designed outside the duplication, which suggests that if these duplications exist, they may not be tandem duplications (which would produce a longer or laddered PCR product) and instead are trans duplications or are segregating within the originally sequenced line. Logically this would fit with the absence of these CNVs in the previous survey which searched for tandem duplications specifically (Chakraborty *et al.* 2017), while dudeML identifies duplications primarily based on coverage and so is agnostic to cis or trans duplications.

Based on these results, bootstrapping appears to average over random effects of simulated training sets to remove a majority of false-positive CNVs called, allowing a more conservative assessment of the copy number variants found throughout an assessed strain. A majority of high confidence false-positives also appear to be actual CNVs, suggesting that dudeML can detect CNVs other tools miss – even using long read data.

## Conclusion

In summary, we have shown that machine learning classifiers, even simple classifiers such as dudeML, perform quite well at detecting copy number variants in comparison to other methods, particularly in samples with reduced coverage or in pools, using just statistics derived from the coverage of a sample. These tools are not computationally intensive and can be used across a large number of datasets to detect duplications and deletions for numerous purposes. We expect machine learning to provide powerful tools for bioinformatic use in the future.

## Acknowledgements

We thank Justin Blumenstiel, Joanne Chapman, Mark Holder, John Kelly, Maria Orive, Daniel Schrider and Carolyn Wessinger for their input in designing the tool, for discussion on machine learning methods and for comments on the manuscript. This work was supported by a K-INBRE postdoctoral grant to TH (NIH Grant P20 GM103418) and by NIH Grants R00 GM114714 and R01 AI139154 to RLU.

## Declarations

### Ethics approval and consent to participate

Not applicable

### Consent for publication

Not applicable

### Authors’ contributions

TH designed dudeML, performed the bioinformatics analysis and statistical analysis, performed the PCR and sequencing, and read and approved the manuscript, RLU designed the CNV detection scheme, provided feedback on tool design, and read and approved the manuscript.

### Data availability

*D. melanogaster* Pacific Bioscience long read data for both Iso-1 and A4 are available on the NCBi short read archive: A4 PacBio SRR7874295 - SRR7874304, Iso-1 PacBio SRR1204085 - SRR1204696. Short read data was downloaded from the following sources: Iso-1 - SRA ERR701706-11, A4 - http://wfitch.bio.uci.edu/~dspr/Data/index.html.

---

**Supplementary Figure 1:**
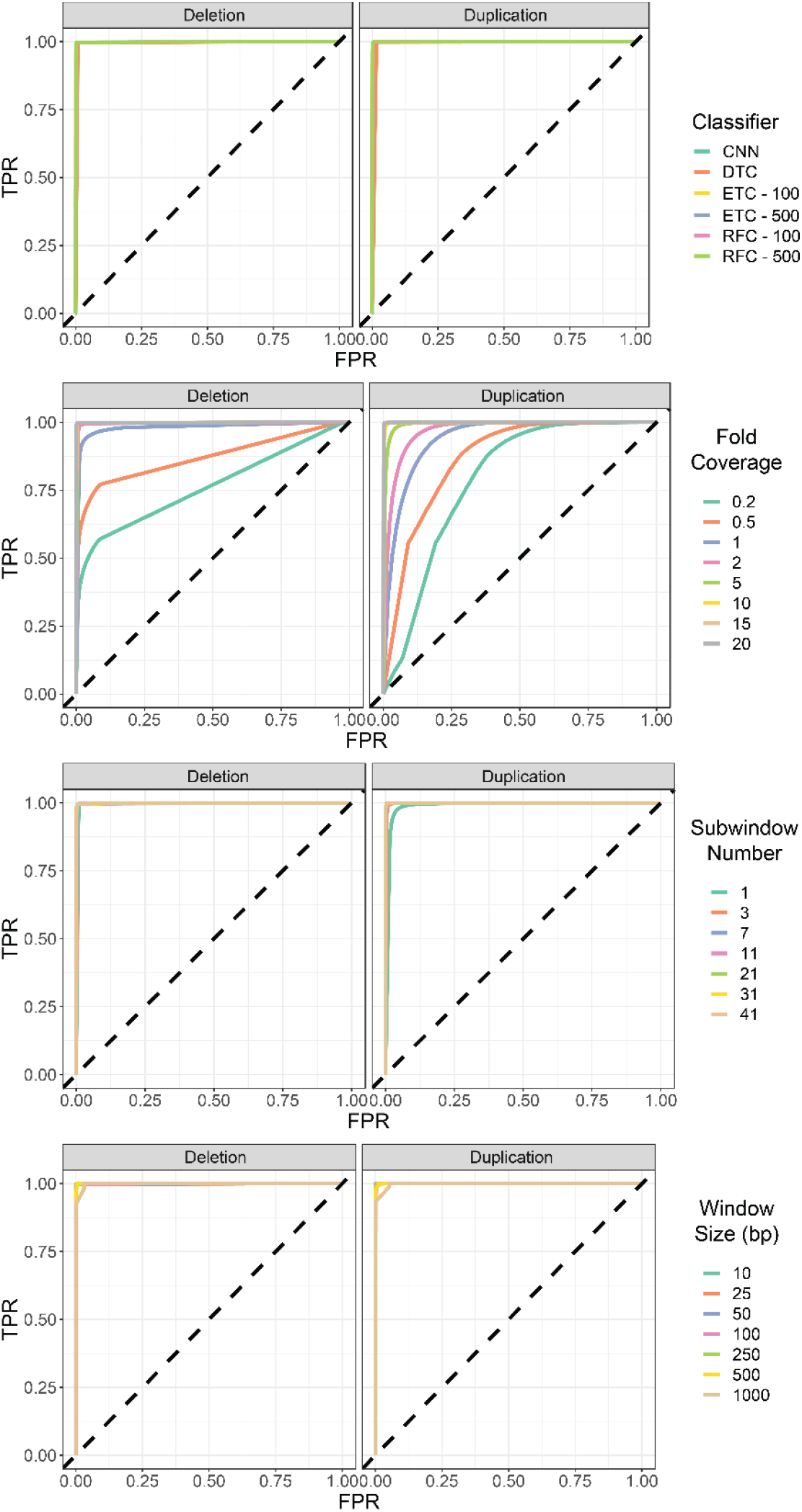
Receiver operating characteristic (ROC) curves for correctly detecting duplications and deletions across different classifiers, sample coverages, sub-window numbers and window-sizes (denoted by line color). Classifiers used as follows: convolutional neural network (CNN), decision tree classifier (DTC), extra trees classifier with 100 estimators (ETC100), extra trees classifier with 500 estimators (ETC500), random forest classifier with 100 estimators (RFC100), random forest classifier with 500 estimators (RFC500).

**Supplementary Figure 2:**
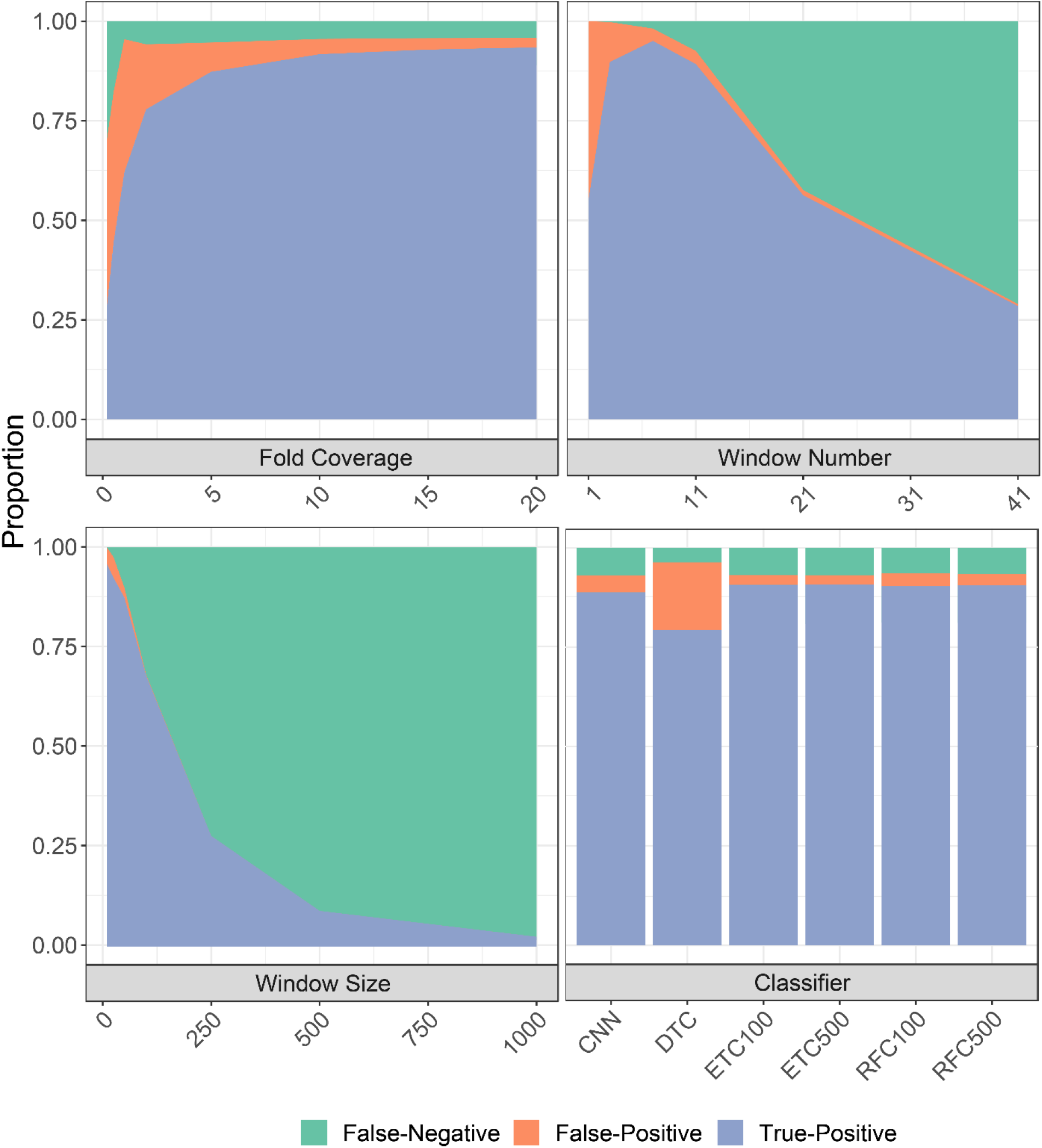
Proportion of CNVs detected or missed given changing parameters, including different numbers of sub windows analyzed, the size of sub windows, the fold coverage of sample data analyzed and different machine learning classifiers used, including convolutional neural network (CNN), decision tree classifier (DTC), extra trees classifier with 100 estimators (ETC100), extra trees classifier with 500 estimators (ETC500), random forest classifier with 100 estimators (RFC100), random forest classifier with 500 estimators (RFC500). If parameter is not variable, it is set as follows: 20-fold coverage, 11 windows, 50bp windows, random forest classifier (100 estimators).

**Supplementary Figure 3:**
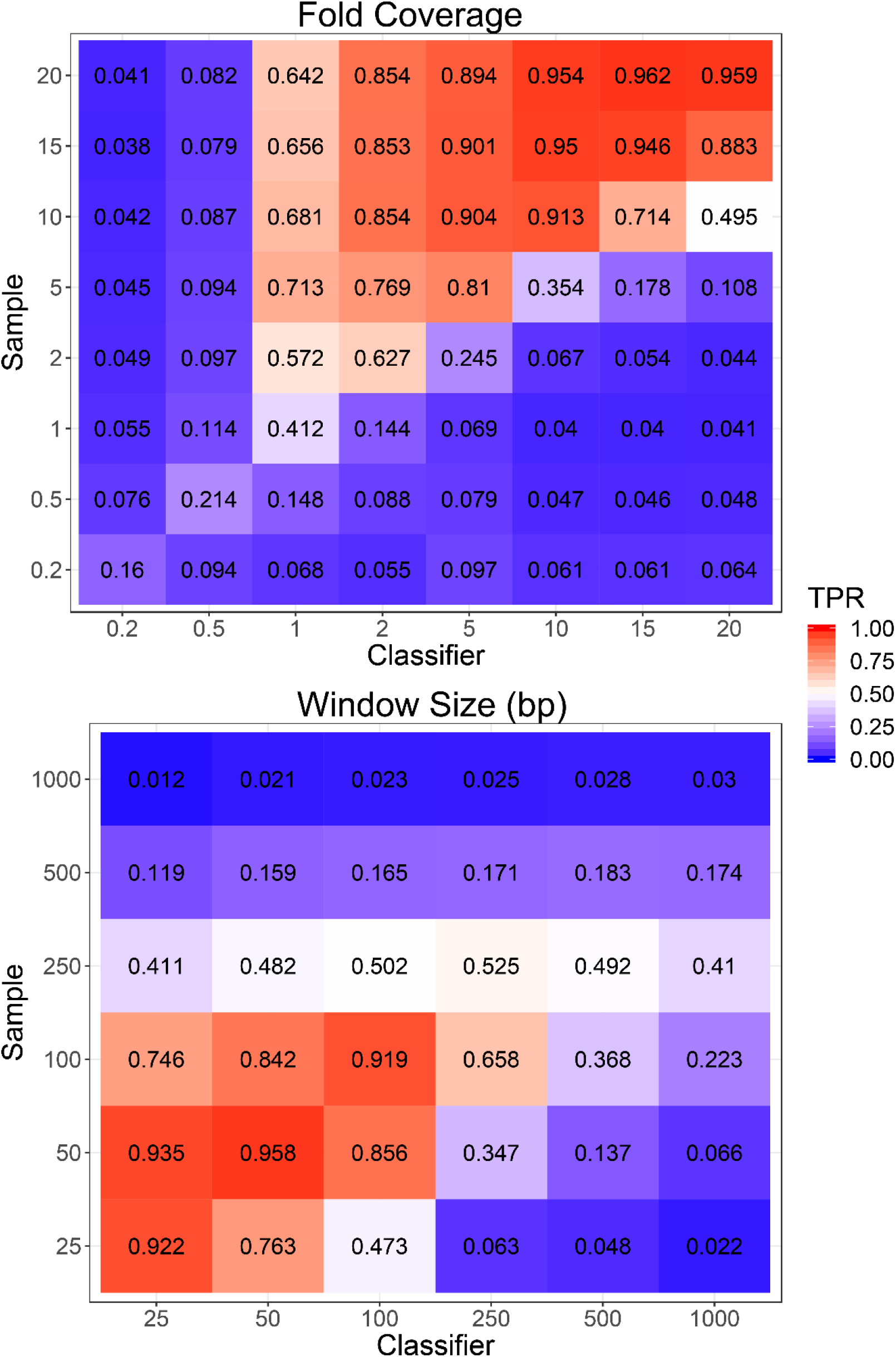
True-Positive rates (TPR) of mis-specified training sets across different fold-coverage samples and classifiers, and different window sizes in samples and classifiers.

**Supplementary Figure 4:**
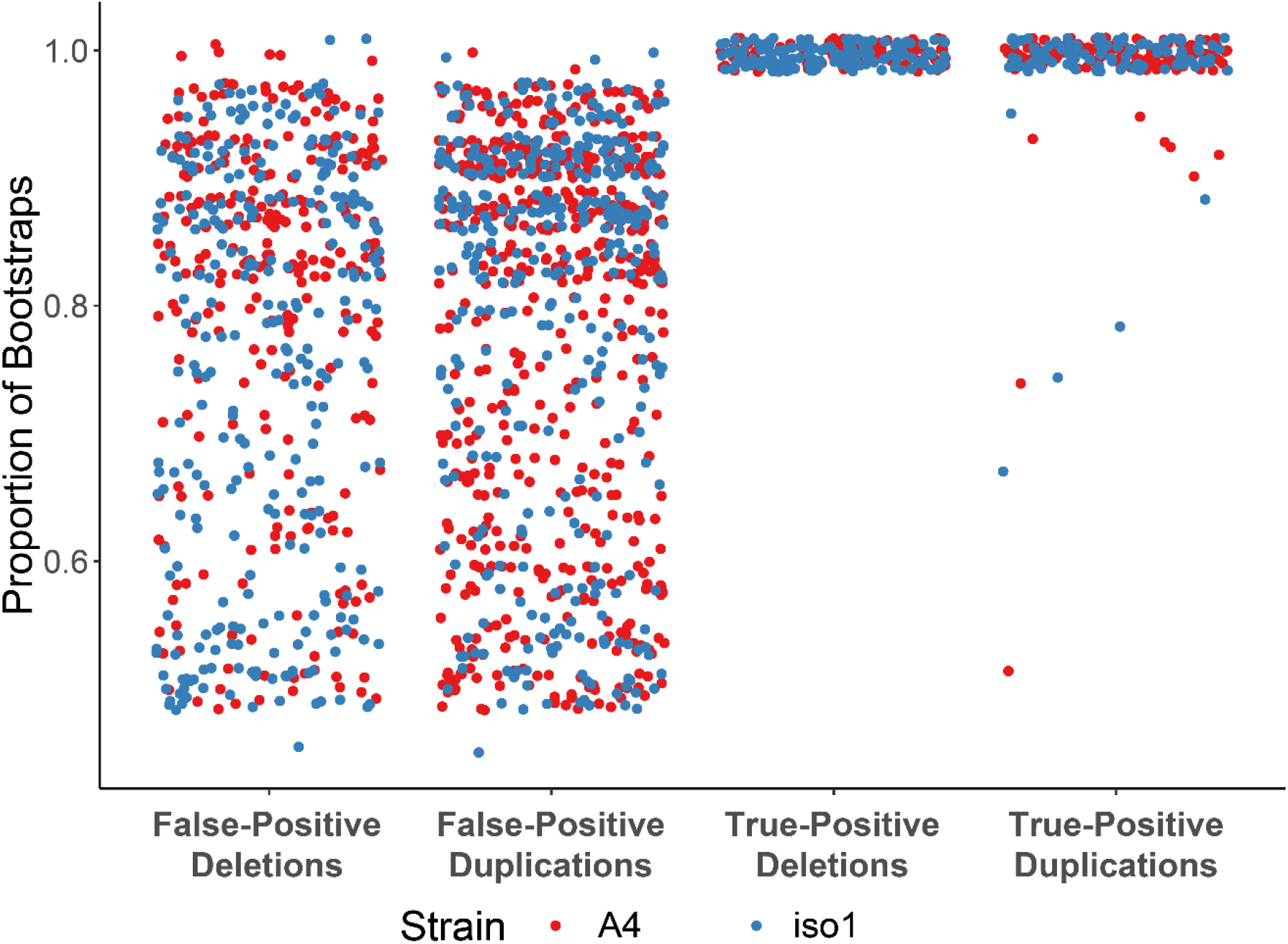
The proportion of bootstraps for each detected CNV in Figure 2, separated by if they are a false-positive, true-positive, duplication or deletion.

**Supplementary Figure 5:**
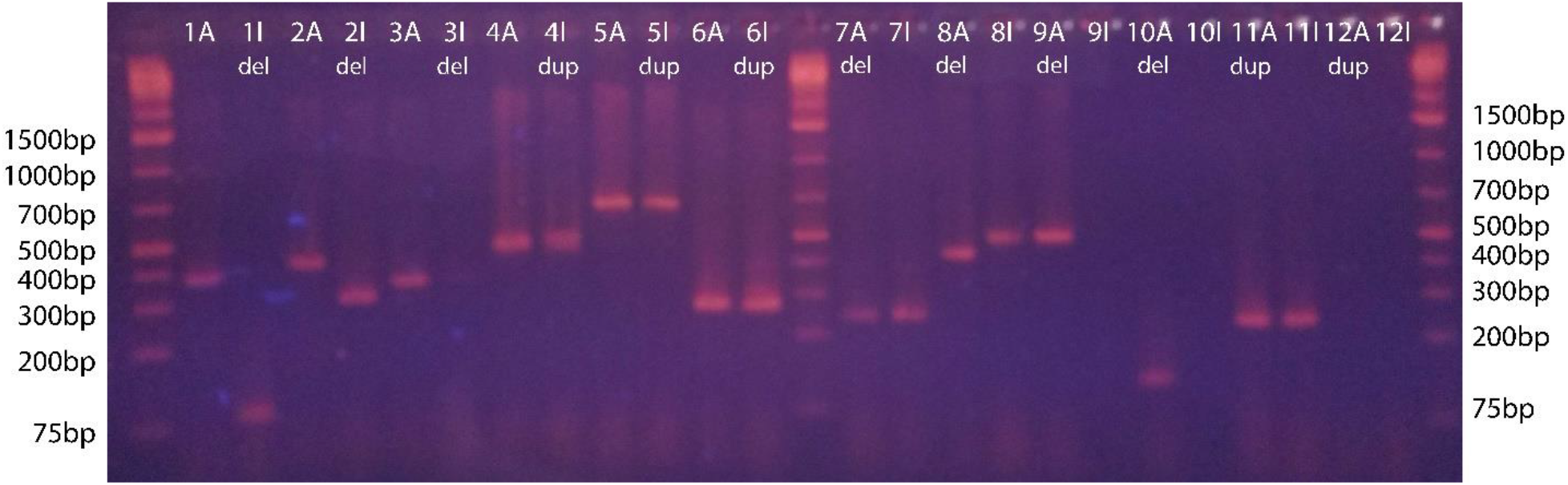
Gel electrophoresis image of PCR products from primers designed around putative CNVs missed in the previous survey, numbered as Supplementary Data 2. Deletions are shown as products shorter than expected, while duplications should be longer or show laddering. Products are ordered showing A4 (A) as the left of the pair, while Iso-1 (I) is on the right.

**Supplementary Data 1:** Screenshots of the integrated genomics viewer for a subset of called duplications and deletions in A4 data mapped to Iso-1 reference genome and vice versa (compared to the data mapped to its own reference). These CNVs were called as false-positives due to their absence in the previous survey. Coverage and reads with supplementary alignments support their existence.

**Supplementary Data 2:** Primer Sequences of a subset of putative duplications and deletions described in Supplementary Figure 5 and Supplementary Data 1.

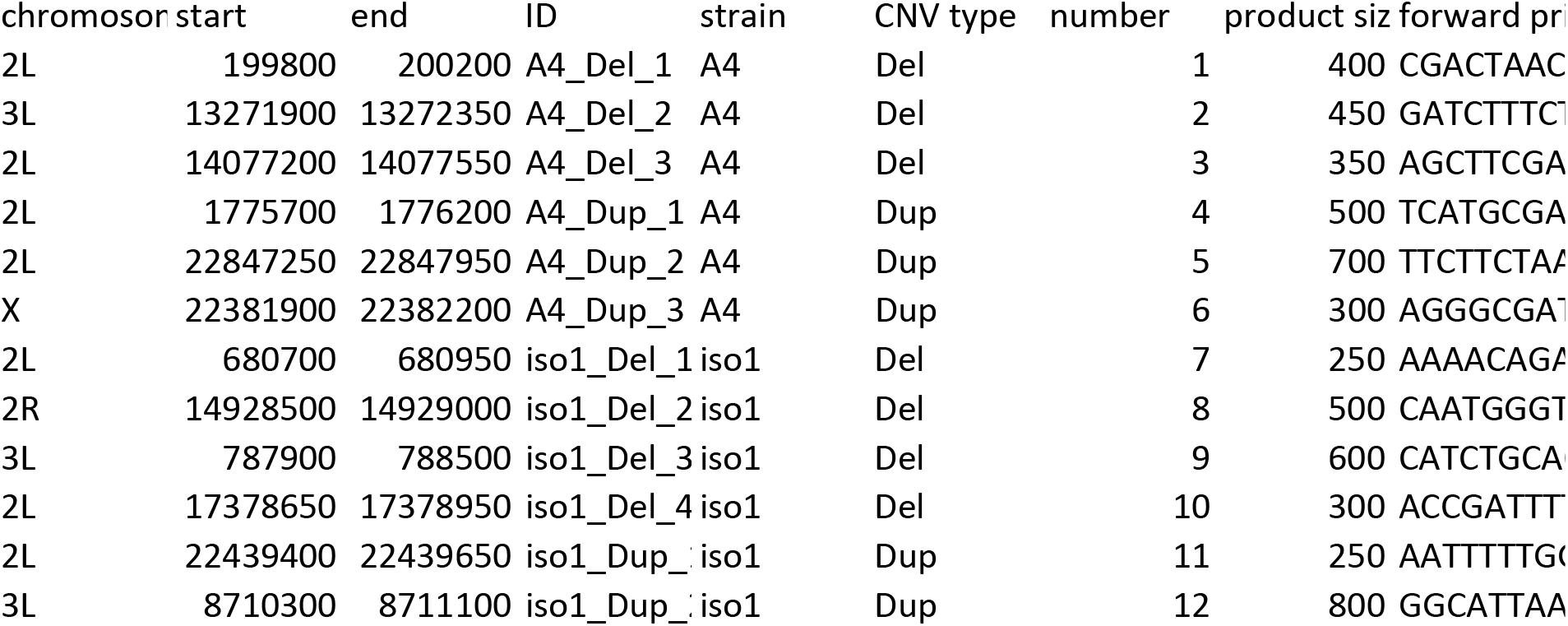
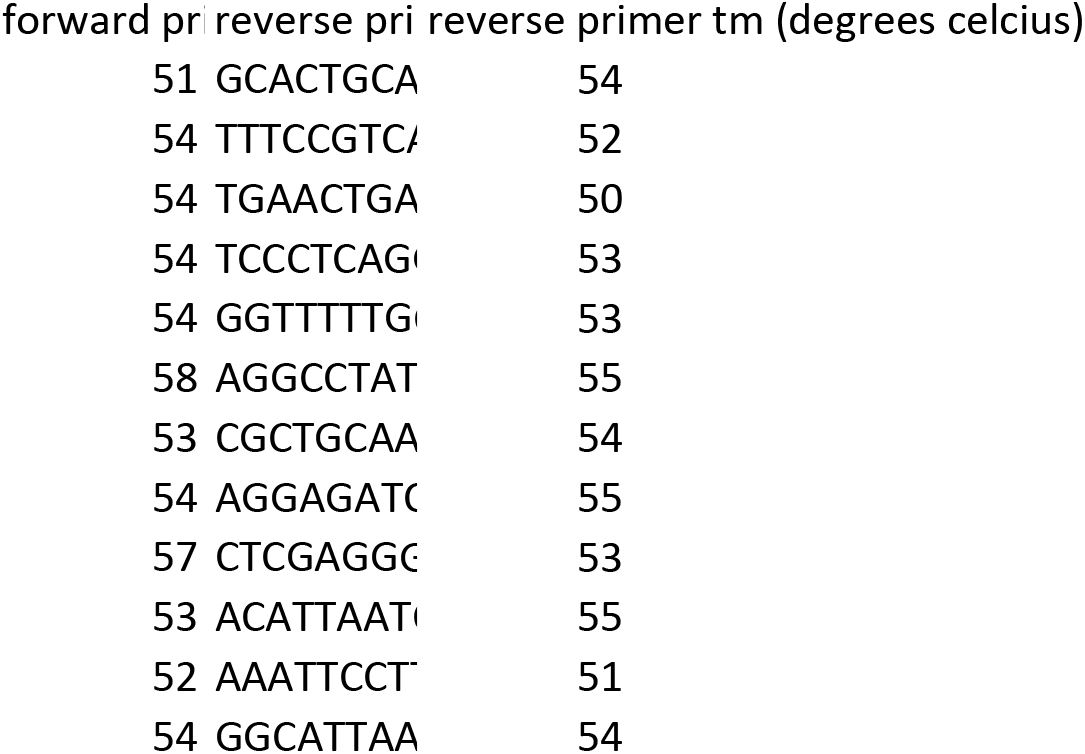

